# *SNCA*^*E46K*^ transgenic *Drosophila* Model of Parkinson’s Disease Confirmed the Causative Role of Oxidative Stress

**DOI:** 10.1101/2020.02.28.969501

**Authors:** Samaneh Reiszadeh Jahromi, S R Ramesh, David I Finkelstein, Mohammad Haddadi

**Author notes:** Correspondence should be addressed to Dr. Mohammad Haddadi, Assistant Professor, Department of Biology, Faculty of Science, University of Zabol, Zabol, Iran.

## Abstract

Parkinson’s disease (PD) is a class of neurodegenerative disorders in which, complex interactions of genetic and environmental agents are involved in the etiology of both sporadic and familial PD cases. α-synuclein-encoding *SNCA* gene is known as one of the major genetic contributors of this disease. *E46K* mutation in *SNCA* gene has not been investigated as intensive as other *SNCA* gene mutations including A30P and A53T. In this study, to induce PD in *Drosophila* flies, *UAS-hSNCA*^*WT*^ and *UAS-hSNCA*^*E46K*^ transgenic fly lines were constructed, where *SNCA* gene was over-expressed in flies brains using GAL4-UAS genetic system. Western blot analysis of head samples of *SNCA*-expressing flies verified *SNCA* expression at protein level. Light and electron microscopy analysis of ommatidial structures were performed to verify neurodegeneration as a result of α-synuclein gene overexpression in *Drosophila* transgenic flies. Confocal microscopy analysis of dopaminergic neuron clusters verified cell loss following *SNCA*^*E46K*^ expression in the flies’ brain. *E46K* α-synuclein gene over-expression resulted in an evident decline in longevity as well as climbing ability of the flies. Biochemical studies of transgenic flies showed a remarkable decline in antioxidant enzymes activity and a significant increase in oxidative markers level as well as AchE enzyme activity. Oxidative stress has been known as a causal factor in PD pathogenesis, following expression of *E46K* mutant version of human *SNCA* gene. This *Drosophila* model is able to facilitate comparative studies of both molecular and cellular assays implicated in the assessment of neurotoxicity of different α-synuclein mutations.

## 1. Introduction

*SNCA* gene is mapped to human chromosome 4q22.1 encoding α-synuclein polypeptide. It was recognized as the primary causative gene in etiology of Parkinson’s disease (PD). Prior to its association with PD, α-synuclein was characterized as a pre-synaptic protein acting at neuron terminals in rat brain (Maroteaux et al., 1988). A human homologue of *SNCA* was discovered in 1993 as the contributor to amyloid plaques deposition in Alzheimer’s disease (AD) (Ueda et al., 1993), The first familial case of PD was described in 1997showing a dominant inheritance pattern for a *SNCA* mutation [A53T] (Polymeropoulos et al., 1997). It was indicated that α-synuclein is a constituent of the lewy body (LB), showing how *SNCA* is completely linked to PD (Xia et al., 2008). Since then, duplication triplications and a number of *SNCA* missense mutations *viz.*, A53T, A30P, *E46K*, Q51D, and H50Q have been associated with Parkinsonism (Kiely et al., 2013). All these mutations are quite uncommon and have highly variable penetrance (Heinzel et al., 2019).

Misfolded proteins and aggregations is a typical feature of lots of neurodegenerative disorders. Lewy body (LB) is the key pathologic characteristic of PD involving α-Synuclein fibrils as its main protein constituent. The involvement of α-synuclein in LB formation may not completely indicate the contribution of this protein to etiology of PD. Nevertheless, mutations in *SNCA* are sufficient to provoke familial PD, with autosomal dominant inheritance pattern.

α-synuclein has been established in a several diverse biochemical forms including membrane-bound alpha helix, β-sheet oligomers, unfolded monomer as well as insoluble LB fibrils (Cookson, 2009). Mutated α-synucleins such as A53T, A30P and *E46K* show a discerning tendency to aggregate. A53T and *E46K* mutants make fibrillar structures very much quicker than wild-type *SNCA* (Conway et al., 2000; Choi et al., 2004).

Oxidative stress (OS) is a well-known causal agent in neurodegenerative process of PD (Litvan et al., 2007). Actually, auto-oxidation of dopamine and the subsequent neuromelanin production in a multifaceted pathway generate reactive oxygen species (ROS) and OS, resulting in dopaminergic neurons failure in PD patients (Hirsch, 1992; Zecca et al., 2001; Sulzer, 2007). Furthermore, higher iron levels in dopaminergic neurons have been contributed to free radicals production through the Fenton-Harber Weiss reaction triggering neurodegeneration process (Dexter et al., 1989b).

*Drosophila melanogaster* is a widely known potent model organism to explore the biology of PD (Feany and Bender, 2000). The above -mentioned statements figured the basis of the present study in order to investigate the neurotoxicity of *E46K* mutation of *SNCA* gene and the implication of OS in transgenic *Drosophila* flies. The GAL4-UAS system was used in order to model PD in the flies. A *UAS-hSNCA*^*E46K*^ transgenic *Drosophila* line was constructed and subsequently crossed to appropriate Gal4 drivers. In addition to the molecular, biochemical, and behavioral assays, microscopic investigations of ommatidial structures were conducted to verify h*SNCA*^*E46K*^ – mediated neurodegeneration and possible causative role of oxidative stress.

## 2. Materials and Methods

### 2.1. Fly stocks

#### Human *SNCA* DNA preparation and cloning

Random integration of P-element transgenesis was applied in order to construct a transgenic *Drosophila* line expressing *hSNCA*, (Southall and Brand 2008). *SNCA* cDNA plasmid (Sinobiological Inc., China) was subcloned into a pUAST vector (Brand and Perimon, 1993). In order to make *SNCA*^*E46K*^, pMT 18T-*SNCA* vector was served as the template for site-directed mutagenesis. DNA segments were sequenced to guarantee sequence accuracy. Afterwards the insert was liberated from positive pUC57 clones and then ligated into pUAST vector. Finally, the *SNCA*^*WT*^ and *SNCA*^*E46K*^ plasmid DNAs were diluted using Milli Q water and were sent to CCAMP Transgenic fly facility, NCBS, Bangalore, India for microinjection into the *w*^*1118*^ *Drosophila* embryos. A total of 5 lines with prosperous injection were achieved for the wild-type and mutated *SNCA* gene, separately. Neuronal expression of the target gene was induced by *elav-Gal4.* Of all the constructed UAS-*hSNCA*^*E46K*^ transgenic fly stocks, the one with the expression level equal to the wild-type *SNCA* was identified using quantitative Western blot analysis. Both the selected UAS-*hSNCA*^*E46K*^ and UAS-*hSNCA*^*WT*^ stocks showed insertion on the second chromosome. In this study, standard genetic markers and 2^nd^ chromosome balancers were used, as well.

The following Bloomington *Drosophila* stocks were used in the current study: *Ddc-Gal4* (# 7010) to induce gene expression in dopaminergic neurons, *elav*^*C155*^ -*Gal4* (# 458) for Pan neural expression, along with *GMR-Gal4* (# 9146) to over express *hSNCA* in *Drosophila* eye structure. *UAS-mCD8::GFP* line (# 32186) was employed to visualize neurons morphology. *w*^*1118*^ (# 5905) was used as the control genotype. All *Drosophila* stocks were raised and kept at 22 ± 1°C, 70-80% relative humidity on standard wheat cream agar media, containing dry yeast granules in a 12 h light/12 h dark cycle in a vivarium. All the study assays were conducted on adult male flies.

### 2.2. Western blotting

This assay was carried out based on the method of Bolt and Mahoney (1997). Total body homogenates of 15 flies of each group underwent standard protein extraction process. The total protein content of each sample was measured. 20 µl of each test sample was mixed with 5 µl of sample loading buffer (6X) and loaded on 12.5% SDS-PAGE. The *w*^*1118*^ stock was served as the negative control. Following electrophoresis, the gel was transferred to the blotting unit, where a nitrocellulose membrane (Cat # LC2000, life technologies, USA) was put on the gel. The unit was filled with Tris-glycine buffer and a 150 mA current was applied for 1.5 h. After protein bands transfer onto the nitrocellulose membrane, the blot was blocked with 3% BSA at 4°C overnight. Then the blot was kept for incubation with 1:5000 anti-human α-synuclein mouse IgG monoclonal antibody (Cat # ABH0261, Invitrogene, USA) (overnight at 4°C). The blot was washed with 0.3 % Phosphate Buffer Saline Triton X (PBSTx) for the duration of 30 min. Next, the blot was incubated with 1:1000 diluted buffer of secondary rabbit anti-mouse HRP conjugated antibody (Cat # 6728, Abcam, UK) in 1X PBS for 1 h at room temperature. The blot was given a wash and afterwards was developed using chromogenic substrate (1X TMB) in order to detect horseradish peroxidase (HRP). The expression of the test protein can be monitored by development of bluish green color bands to which secondary antibody is bound. Anti β-actin antibody (Cat # 8224, Abcam, UK) was utilized as loading control.

### 2.3. Scanning electron microscopy (SEM)

SEM was conducted to investigate surface morphology of *E46K* transgenic flies eyes using the method of Kimmel et al., (1990). Adult flies were anesthetized and preserved in 25% ethanol for the duration of 24 h at room temperature. Afterwards, the samples were dehydrated in 50%, 75% and 2 × 100% ethanol for 24 h at each step. Furthermore, the samples got dried using hexamethyldisilazane (HMDS). Then to allow HMDS to evaporate, uncapped vials were kept under fume hood overnight. Dried samples were examined under Carl Zeiss EVO LS-10 SEM.

### 2.4. *Drosophila* eye preparation for ultra-microtomy

Preparation of adult *Drosophila* eye for thin sectioning was performed based on Mishra and Knust (2013) procedure, with a few slight changes. Heads of adult *Drosophila* flies were dissected out and subsequently fixed in the mixture of 2.5% glutaraldehyde and 2% paraformaldehyde in PB at 4°C overnight. Proboscis structures were removed to improve the penetration of the fixative solution. Heads were washed with PBS (three times, with the time intervals of 10 min) and incubated in 0.5% osmium tetroxide as the secondary fixative for 2 h in dark condition at room temperature. Next, samples were washed with distilled water (three times, with the time intervals of 10 min). Samples dehydration was carried out by serial ethanol treatment (50%, 70%, 90%, 95%, and 2 × 100%) for 10 min, and subsequent two washing stages with propylene oxide for 10 min. The infiltration was achieved by storing test tissues in a solution mixture of fresh epoxy resin as well as propylene oxide while holding on a shaker at the room temperature. To embed the samples, flat-embedding molds were filled with freshly prepared epoxy resin and after that samples were placed within the blocks, as well. The orientation of the samples were aligned properly t under a stereomicroscope. To achieve polymerization, blocks were incubated at 60° C for the duration of 48 h. Plastic blocks were cut into the sections of 1µm thickness using Leica EM UC6 ultramicrotome (Leica Mikrosysteme, Austria). Semi-thin sections were placed on a slide and got dried on a slide warmer tool following1 min staining with Toluidine blue. Sections were washed, dried and mounted using DPX (Leica Mikrosysteme, Austria) and finally observed under light microscope.

### 2.5. Confocal microscopy

To inspect and assess CNS dopaminergic neurons in *E46K* transgenic Drosophila flies, brain dissection was carried out on ice and subsequently handled for fixation using cold 4% paraformaldehyde for the duration of 30 min. Then, PBSTx was used to wash brain samples for 40 min, with 4 changes per 10 min using a shaker at the room temperature. At last, samples were mounted using Vectashield mounting medium (Vector Laboratories, USA). The prepared samples were examined by confocal microscope (LSM 710, Carl Zeiss, Germany).

### 2.6. Survival assay

To assess the *SNCA* neurotoxicity survival rate of flies lately eclosed flies were clustered in groups of 20. The flies were transferred to fresh media in 3-days intervals. The number of dead flies was quantified in each turn over and alive insects were considered for calculation of survivorship (Haddadi et al., 2016a). Based on the observed longevity among the transgenic flies, 20-day old adult flies were used for further behavioural and biochemical investigations.

### 2.7. Negative geotaxis assay

The locomotory activity of the flies was characterized by negative geotaxis assay as formerly described (Jahromi et al., 2013). Thirty adult male flies were placed in a scaled plastic tube. Flies were kept for 10 min rest to acclimatize to the new environment. Next the flies were slightly tapped down and then were let to climb up. The number of flies that crossed the 20 cm mark on the tube in 1 min was quantified. Six biological replicates of each genotype were analyzed and the corresponding values represented as mean ± standard error.

### 2.8. Locomotion tracing assay

To trace locomotion behaviour of the flies the iFly system was employed (Kohlhoff et al., 2011). This assay analyses the flies’ movement in an automated fashion. The iFly uses a single digital camera to track the trajectories of up to 20 individual flies. Briefly, ten flies were transferred to a glass tube and then placed in a chamber that is covered by a frosted-plastic lid. To facilitate 3-D tracking, two mirrors were placed at equal angles at the back of the tube to allow side images capture by the camera. A 10-min rest was given to the flies for acclimatization to the new place. Next, flies were gently tapped down every 30 second and their locomotion was recorded for 90 seconds. The captured video clips were analyzed by C-trax software (Kohlhoff et al., 2011). Movement of each individual fly was quantified and corresponding graphs were plotted by the software.

### 2.9. Ethanol exposure

The standard protocol of Maples and Rothenfluh (2011) was conformed to measure ethanol sensitivity of the flies. For this assay, eight flies were collected one day before ethanol exposure. The ST50 (the time required to get half of the flies stationary) and RC50 (the time required for complete recovery of half of the sedated flies) were measured and expressed in minutes. ST50 and RC50 values of eight biological replicates of each genotype were averaged and represented as mean ± standard error.

### 2.10. Biochemical bioassays

The samples for biochemical assays were homogenates of heads of 100 flies suspended in ice-cold phosphate buffer (PB) and centrifuged at 3000*g* for 10 min at 4°C. The supernatant was subsequently utilized to assess activity of antioxidant enzymes and levels of oxidative markers.

As we previously described (Haddadi et al., 2016b) the catalase (CAT) activity was assessed based on the H_2_O_2_ hydrolysis rate (Aebi, 1984). The enzyme activity was expressed as µM of H_2_O_2_ used/min/mg protein. Superoxide dismutase (SOD) activity was measured based on SOD-mediated inhibition of pyrogallol autoxidation (Marklund and Marklund, 1974). The enzyme activity was shown as units; wherein 1 unit corresponds to 50% inhibition of pyrogallol auto-oxidation. Acetylcholinesterase (AChE) activity was assessed as per Ellman et al., (1961) by employing acetylthiocholine (a sulfur analogue of acetylcholine). After hydrolysis, acetylthiocholine generates acetate and thiocholine. Thiocholine makes a yellowish compound (5-thio-2-nitrobenzoate anion) in presence of DTNB. The enzyme activity was shown as nmoles of DTNB hydrolyzed/min/mg protein. Reduced glutathione content was assessed according to the method of Hissin (1976) using O-phthalaldehyde (OPA). In the assay, OPA reacts with GSH and generates strong fluorescence signals that can be used for estimation of GSH composition. The glutathione level was estimated using a standard curve and was represented as µg GSH/mg protein. ROS levels were measured based on the fluorometric method with DCFH-DA probe (Black and Brandt, 1974). If ROS are present in reaction, the non-fluorescent DCFH-DA probe gets quickly oxidized into 2′, 7′-dichlorofluorescein (DCF), a highly fluorescent agent that can be spot using fluorometric measurements. The DCF concentration in test samples was calculated using DCFH-DA standard curve. Lipid peroxidation (LPO) was assessed as per Ohkawa et al. (1979) using thiobarbituric acid (TBA). The biochemical basis of LPO assay is malondialdehyde (MDA) synthesis which is an end-product of LPO process which reacts with TBA and form a chromogenic solution. The MDA contents of samples was measured using tetramethoxypropane molar extinction coefficient (ε) value which is equal to 15600 M^-1^cm^-1,^ the. Estimation of total protein content was performed according to Lowry et al., (1951). The method is based on the reaction of peptide nitrogen and the copper ions in alkaline condition and consequent Folin-Ciocalteu reagent reduction through copper-catalyzed oxidation of aromatic acids. The protein content of each sample was assessed using bovine serum albumin (BSA) standard curve.

### 2.11. Statistical Analysis

Data was presented as mean ± SE. Mean comparison between groups was carried out by independent *t*-test in SPAW (predictive analytics software) version 19.0. *p* value of 0.05 was assumed as the minimum level of significance. Three significance level was considered, each represented by different asterisks numbers as demonstrated in all illustrations (\**p* < 0.05, \*\**p* < 0.01 and \*\*\**p* < 0.001).

## 3. Results

In the present study, *E46K* α-synuclein transgenic *Drosophila* stock was constructed and Gal4-mediated α-synuclein gene expression in transgenic flies’ brains was verified at the protein level. In this regard, western blot analysis on *E46K;elav* transgenic flies head samples was carried out. The results confirmed successful Gal4-UAS mediated over-expression of *E46K* mutant form of human α-synuclein gene in transgenic flies (Fig. 1). Following confirmation of *E46K* α-synuclein protein expression, a critical assessment was performed to test the ability of *E46K* form of α-synuclein protein in inducing neurodegeneration process in transgenic flies. Hence, in the first stage, *E46K* α-synuclein gene over-expression in *Drosophila* eye structure was carried out by means of *GMR*-Gal4 driver; following which eye structures of 10 day-old F_1_ progeny were observed under a stereo microscope. The results of SEM studies in these transgenic lines showed that *E46K* α-synuclein gene over-expression in retinal structures results in degeneration of retinal neurons, a process which was obvious by observing irregular eye surface morphology (Fig. 2). Light microscopy analysis of the apical tangential sections of the fly ommatidia (Fig. 3) revealed disorganized photoreceptor cells within the omamatidial structure and a higher level of neurodegeneration, evident by the observation of vacuoles in *E46K* α-synuclein transgenic flies. *Ddc*-Gal4 expression pattern was further assessed using *Ddc*-GFP reporter gene complex. Our results showed that Ddc expression pattern includes dopaminergic neurons of adult *Drosophila* brain. Confocal images of the brain of *E46K* α-synuclein transgenic flies reveal neurodegeneration in protocerebral antero-medial (PAM), protocerebral posterio-medial (PPM) as well as protocerebral posterio-lateral (PPL) clusters of dopaminergic neurons (Fig. 4).

**Figure 1.**
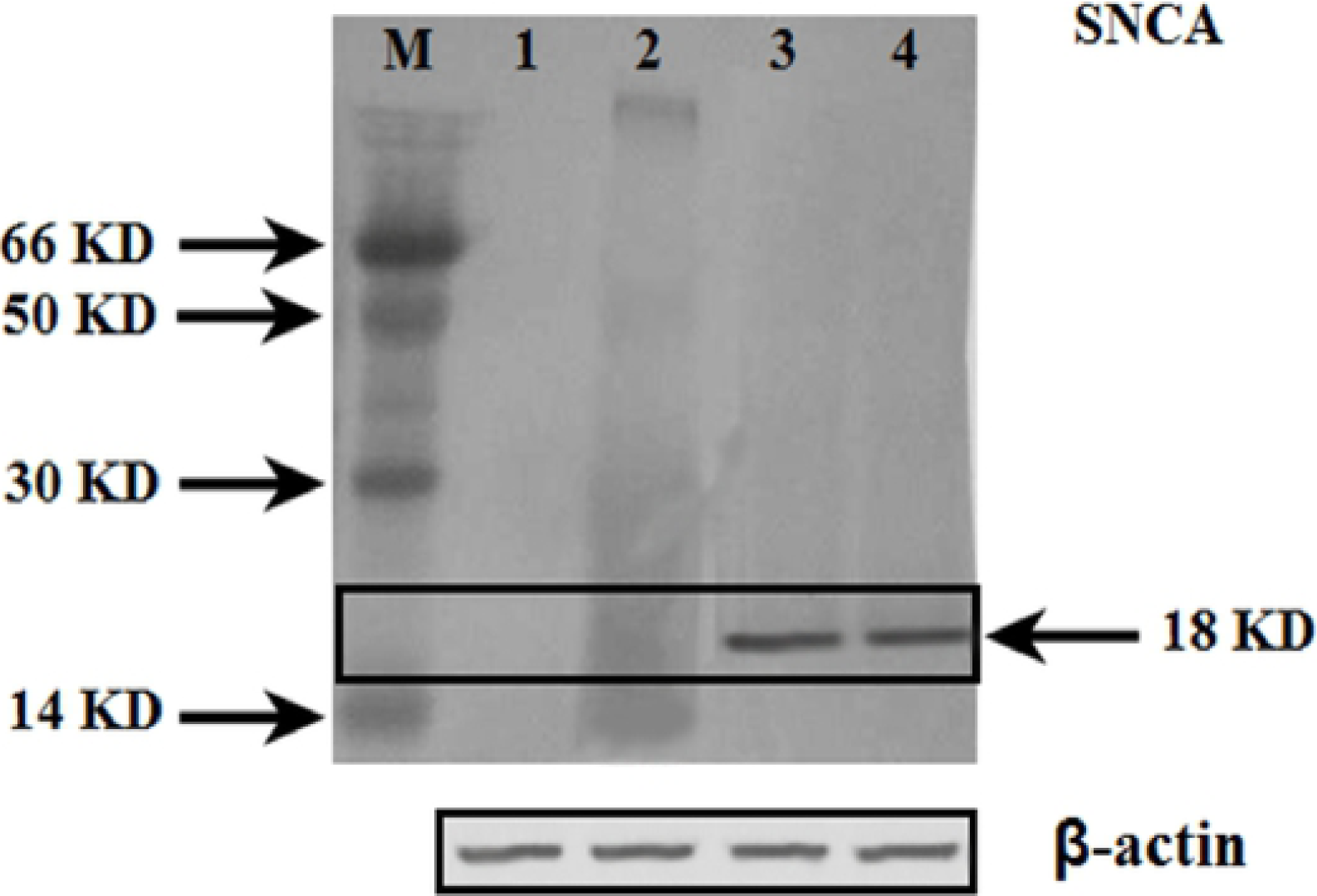

**Figure 2.**
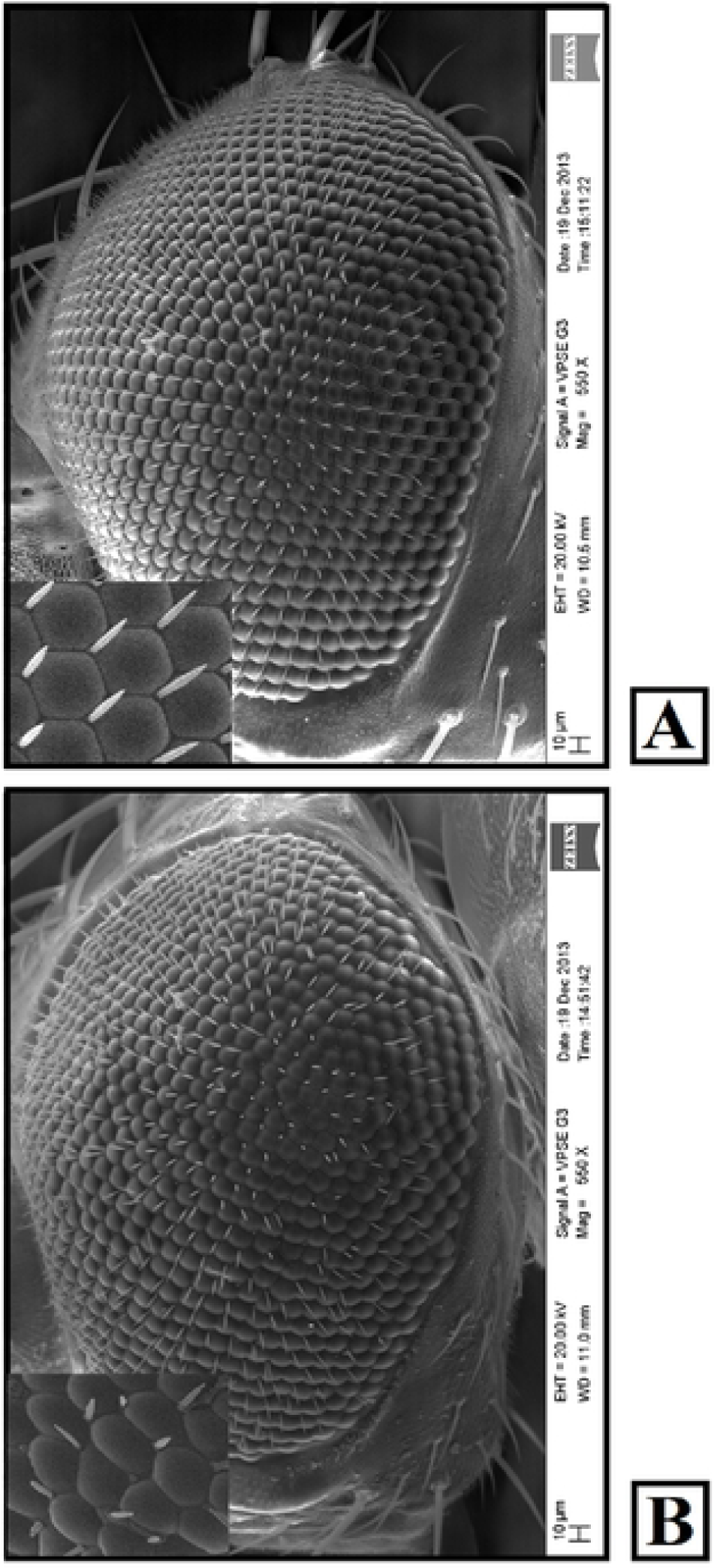

**Figure 3.**
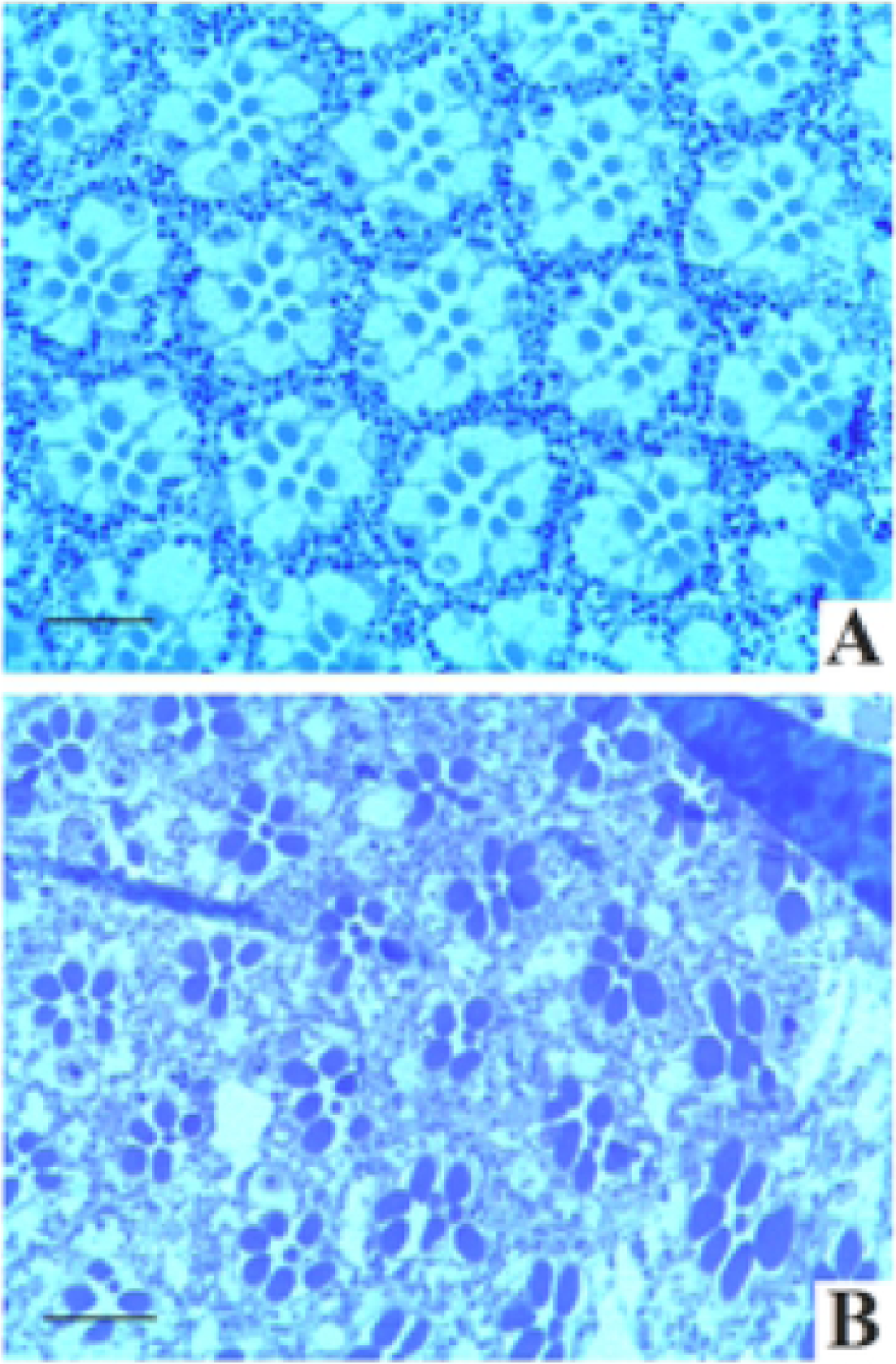

**Figure 4.**
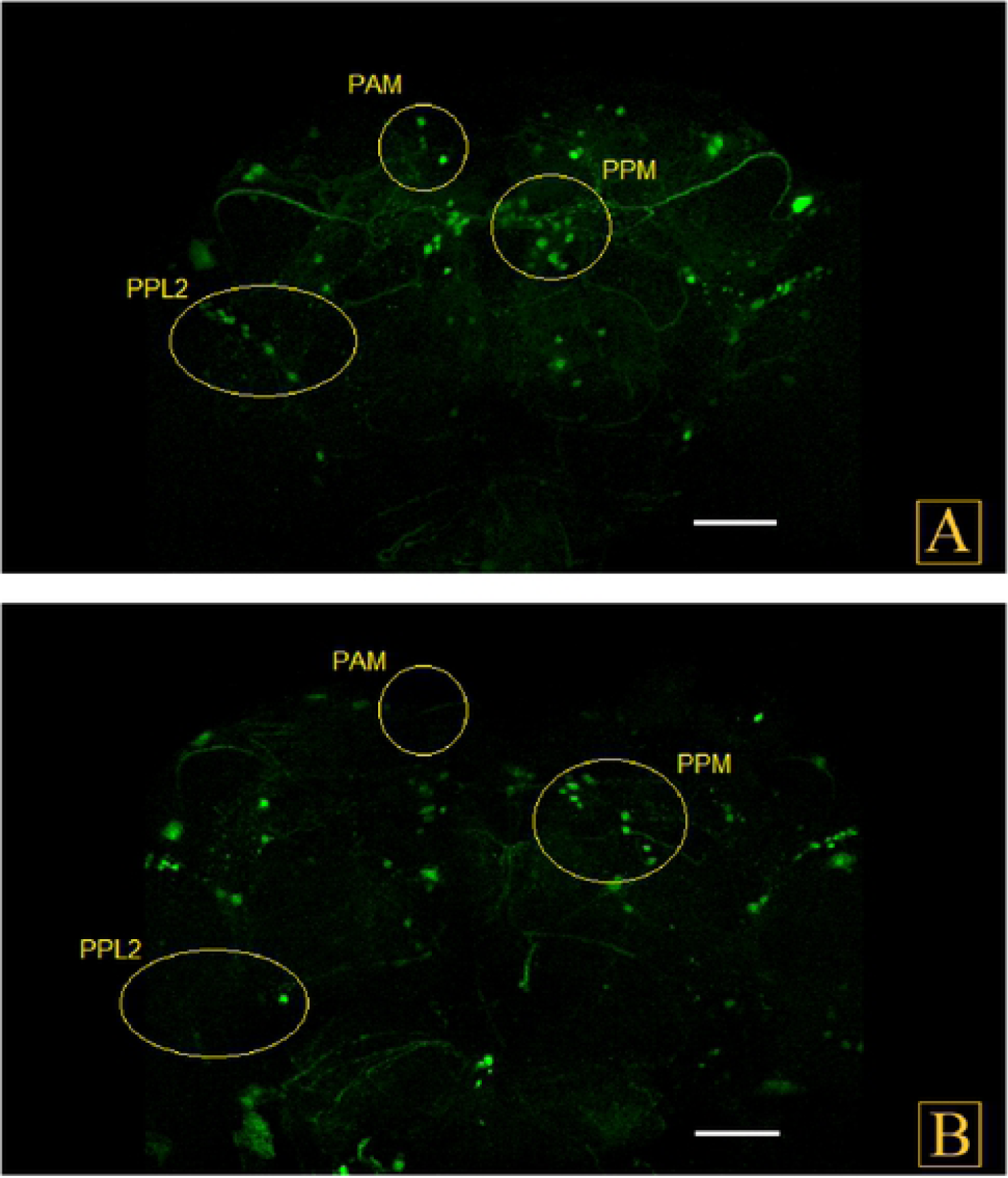

After confirmation of over-expression of *E46K* mutant type α-synuclein gene in the brain of transgenic flies and the consequent α-synuclein-mediated neurodegeneration, behavioral and biochemical assessments were undertaken to validate previous observations.

Survival assay illustrate that PAN neuronal over expression of *SNCA* led to reduced lifespan of the transgenic flies (Table 1.). Maximum lifespan of *w*^*1118*^ flies was 90±3 days while the same parameter was found to be 81±1 and 75±2 days for *UAS-SNCA*^*WT*^*/+; elav/+* and *UAS-SNCA*^*E46K*^*/+; elav/+* genotypes, respectively. The mean lifespan of *SNCA* expressing flies displayed a significanr reduction when compared to the control genotype (*p* < 0.001, *n* = 360).

**Table 1:**
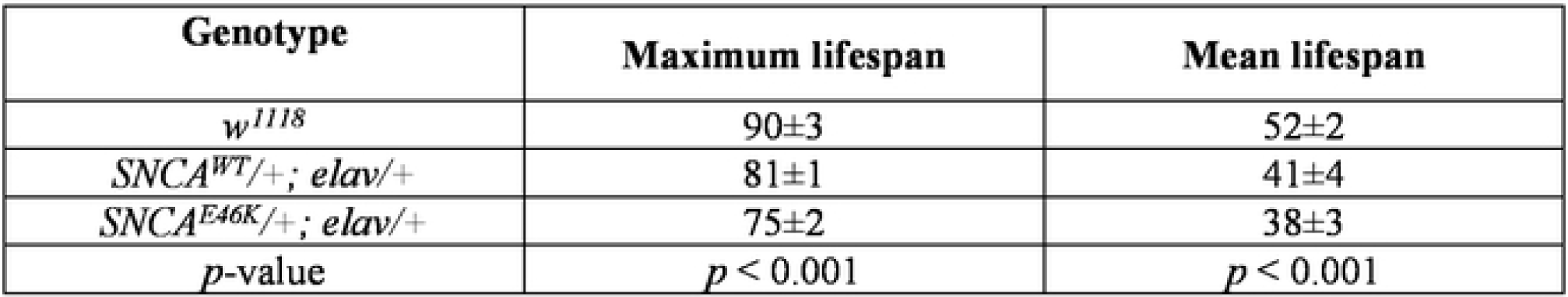
Mean±SE of lifesapn in normal and *SNCA* transgenic flies. *SNCA*^*E46K*^ expressing PD model flies live shorter and have lower longevity rate. At least 360 one-day old adult flies of each genotype were assayed for longevity. Statistical analysis revealed significant decline in maximum and mean life span of the PD transgenic insects (*p* < 0.001).

Detailed observation of Figure 5 clearly showed that the climbing ability of transgenic flies in negative geotaxis assay is diminished as compared to control ones. Automated tracking of individual flies indicate lower movement rate of SNCA^E46K^ model flies and also it was shown that these flies pretend to stay at the bottom of test tubes while the control flies travel faster and climb up to the middle and top parts of the tube (Fig. 6).

**Figure 5.**
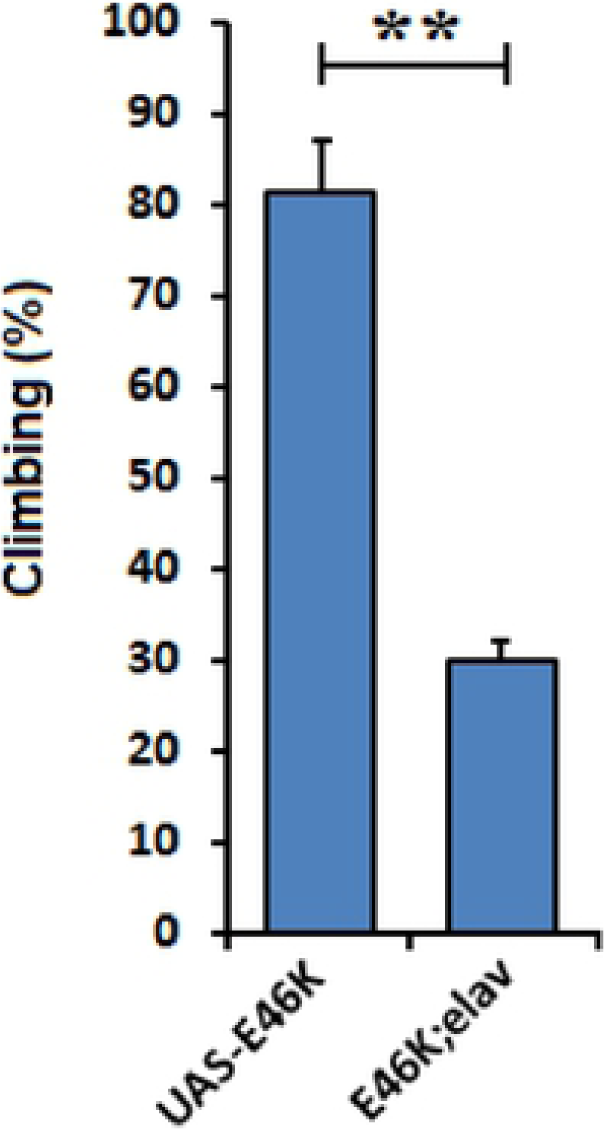

**Figure 6.**
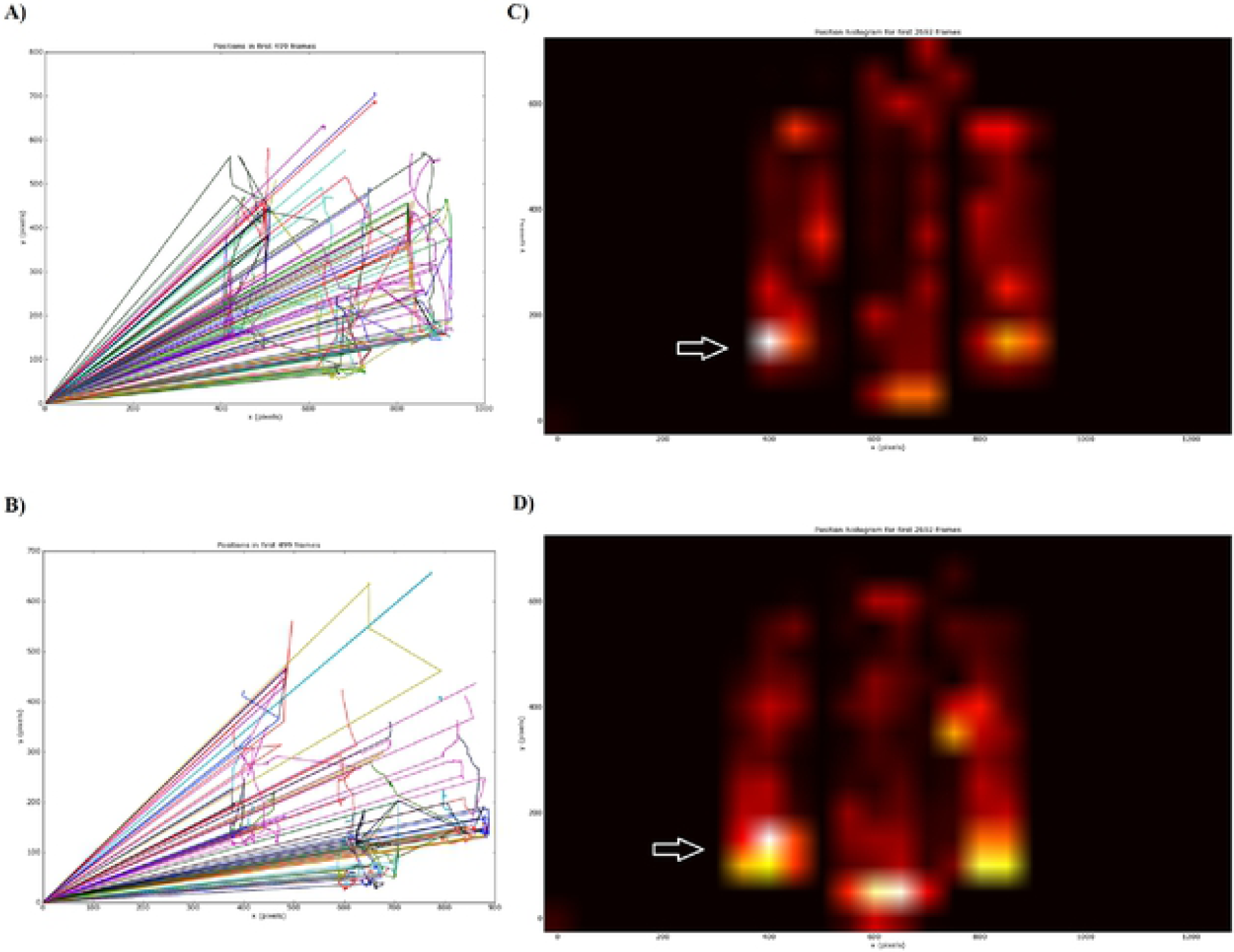

Behavioural assays regarding ethanol sensitivity were prepared by valuation of ST50 and RC50 scores in the transgenic flies. The subsequent results are represented in Fig. 7. A perusal of the figure indicates higher sensitivity and lower resistant to ethanol vapour among E46K α-synuclein expressing flies which was evident by lower ST50 and higher RC50 rates compared to the control genotype.

**Figure 7.**
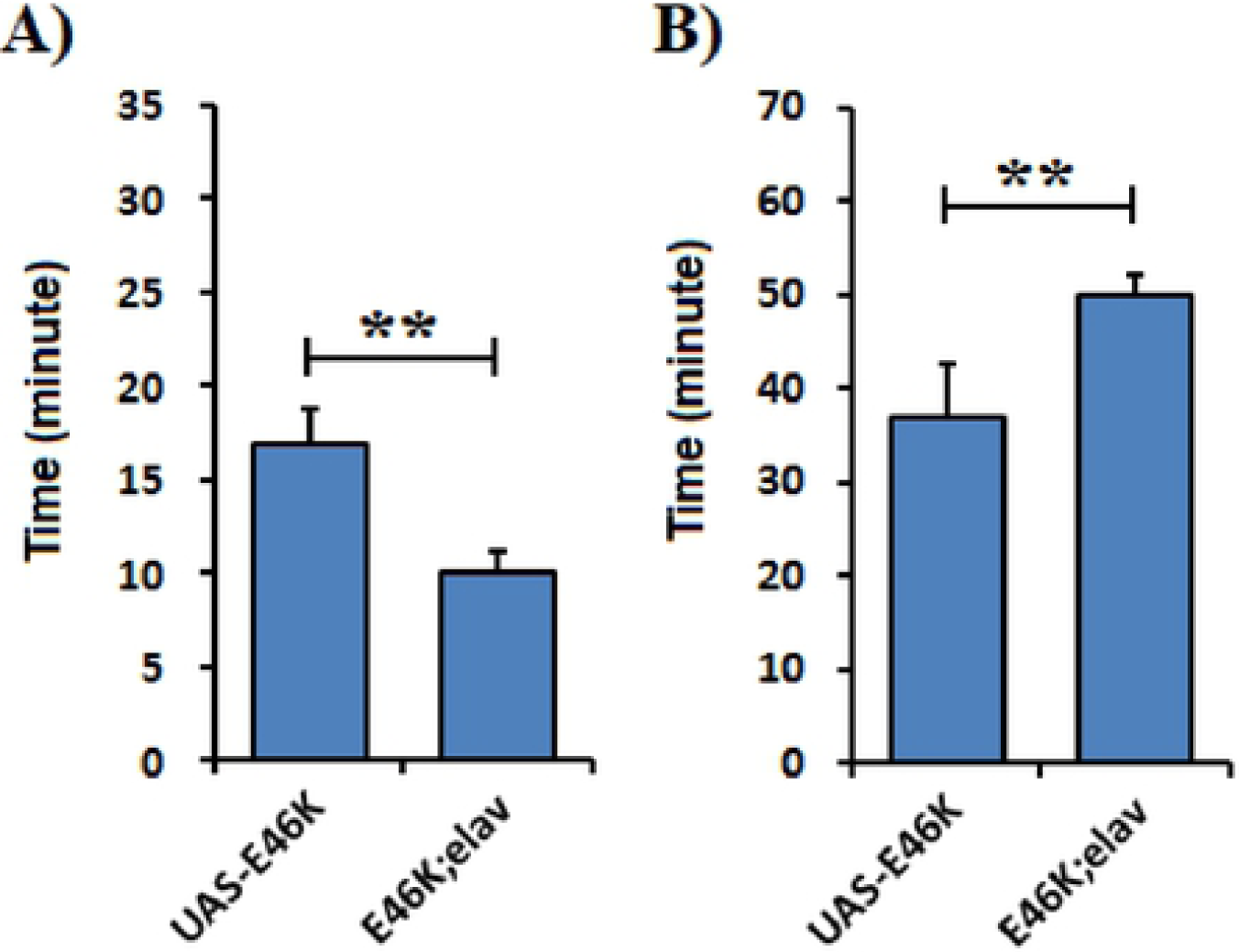

The results on biochemical assessment of enzymes activities and OS markers levels are compiled in Table 2. The results revealed that *E46K* α-synuclein gene over-expression induces decreased CAT and SOD activities in transgenic flies as compared to controls. CAT and SOD activities were measured as 4.4±0.53 μM H_2_O_2_/min/mg protein and 0.47±0.03 unit/mg protein, respectively, in transgenic flies and 6.5±0.27 μM H_2_O_2_/min/mg protein and 0.62±0.04 unit/mg protein,, respectively, in control ones (*p* < 0.01, *n* = 4). LPO and ROS levels were significantly higher in transgenic flies than control ones. LPO and ROS levels in transgenic flies were measured as 6.8±0.26 nM MDA/mg protein and 895±21.6 nM DCF/mg protein, respectively. The results revealed significant depletion in GSH level (12.8±1.2 μg GSH/mg protein) in transgenic flies as compared to controls (18.5±1.34 μg GSH/mg protein) (*p* < 0.01, *n* = 4). While AchE activity was measured as 145±3.6 μM/min/mg protein in transgenic flies; it was found to be 92±8.4 μM/min/mg protein in control ones (*p* < 0.01, *n* = 4).

**Table 2:** Msean±SE of endogenous antioxidant enzyme activity, oxidative markers and AchE in *SNCA* transgenic flies.

| Genotype | AchE | CAT | SOD | GSH | LPO | ROS |
| --- | --- | --- | --- | --- | --- | --- |
| *UAS-SNCA<sup>E46K</sup>* | 92.0±8.4 | 6.5±0.27 | 0.62±0.04 | 18.5±1.34 | 5.0±0.47 | 700±33.1 |
| *Elav; SNCA<sup>E46K</sup>* | 145.0±3.6 | 4.4±0.53 | 0.47±0.03 | 12.8±1.02 | 6.8±0.26 | 895±21.6 |
| *p*-value | *p* < 0.01 | *p* < 0.01 | *p* < 0.01 | *p* < 0.01 | *p* < 0.01 | *p* < 0.01 |

AchE = Acethylcholinesterase CAT = Catalase specific activity (μM H<sub>2</sub>O<sub>2</sub>/min/mg protein) SOD = Superoxide dismutase specific activity (units/mg protein) GSH = Glutathione levels (μg GSH/mg protein) LPO = Lipid peroxidation levels (nM MDA/mg protein) ROS = Reactive oxygen species levels (μM DCF/mg protein)

## 4. Discussion

PD is a multifactorial disorder apparently possessing a polygenic inheritance pattern. Exposure to some of the environmental neurotoxins, and gene-environmental interactions are the factors being considered as important in the etiology of this disease (Bonilla-Ramirez et al., 2013). Though most of the PD cases are idiopathic (no known cause) autosomal recessive parkinsonism or autosomal dominant parkinsonism all share synuclein deposition (Bekris et al., 2010). *SNCA* is one of these related genes in which missense mutations (A30P, A53T, *E46K*, G51D, H50Q) cause familial form of PD (Si et al., 2017). Therefore, α-synuclein appears to be a crucial gene in both familial and sporadic forms of PD.

The mutations in human are rare, the penetrance, pathobiology, and biochemistry have not been fully investigated (Heinzel et al., 2019). Genetic manipulation methods such as making transgenic *Drosophila* are valuable tools to assess genetic malfunctions in human neurodegenerative disorders. Such genetic malfunctions can be achieved by over-expression of selected genes under the control of UAS enhancer sequence in binary combination with GAL4 as a transcription factor that can restrict the expression of gene of interest both temporally as well as spatially (Brand and Perrimon, 1993). In this view, in the present study as an additional approach to model PD in *Drosophila*, transgenic flies bearing *E46K* mutant form of *SNCA* gene were constructed, where *Drosophila* CNS neurons were targeted for α-synuclein gene over expression. In addition, cellular antioxidant defense system, neurotransmitter enzyme profile and *E46K* α-synuclein transgenic flies behavioral impairments were evaluated to address the issue of neurodegeneration.

Ever since most of fly PD-like symptoms in α-synuclein transgenics are detected after day 10-12. Assessment of stable age-related molecular modification occurring prior to the beginning of neurodegeneration could reveal the casual events in the pathology of PD (Feany and Bender, 2000; Xun et al., 2007). *SNCA*^*E46K*^ transgenic flies used in the present study demonstrates the most significant behavioral age-dependent defects in their climbing capability as compared to the control ones.

The results of our assessments in *Drosophila* verified the association of α-synuclein over expression with striking movement defects as characterized by negative geotaxis assay. Our results are in agreement with the results of other studies on locomotor function of α-synuclein transgenic *Drosophila* and mice (Wassef et al., 2007; Kudo et al., 2011b).

The role of oxidative stress in the etiology of neurological diseases has been implicated (Lin and Beal, 2006). Oxidative stress is thought to be a series of pivotal biochemical events that may causes the onset of PD, α-synuclein aggregation, and dopaminergic neurons degeneration (Quilty et al., 2006). Intra-cellular over expression of α-synuclein and the consequent ROS is involved in plasma membrane damage, mitochondrial dysfunction, and decline in glutathione level, all of which make the brain susceptible to oxidative damage (Mancuso et al., 2007).

Our results illustrated significant elevation of ROS and LPO level and marked depletion of glutathione level in α-synuclein-expressing transgenic flies. Our findings are in line with those of other studies demonstrating higher levels of ROS as well as LPO in PD model of *Drosophil*a (Casani et al., 2013; Siddique et al., 2013). Interestingly, our results demonstrated diminished activity of CAT and SOD enzymes in *E46K* α-synuclein-expressing flies, as was reported in DJ-1 mutant flies (Casani et al., 2013) and *SNCA* A30P flies (Jahromi et al., 2013), as well. Furthermore, *E46K* α-synuclein flies showed an increased AchE enzyme activity, indicating the neurotoxicity in the brain of transgenic flies.

The present study provided a powerful evidence for contribution of oxidative damage in the onset of PD symptoms in *SNCA*^*E46K*^ transgenic flies. Although the same scenario is applied to other *SNCA* mutations, gradual and age-related neurodegenerative process in PD fly models makes systematic and comparative assessments possible, targeting efficient disease management strategies, prior to the start of its visible symptoms.

## Acknowledgements

This research project was partially supported by Iran National Institute for Medical Research Development, grant No. NIMAD-957644.

